# Piezo1 balances membrane tension and cortical contractility to stabilize intercellular junctions and maintain epithelial barrier integrity

**DOI:** 10.1101/2024.11.20.624378

**Authors:** Ahsan Javed, Aki Stubb, Clémentine Villeneuve, Franziska Peters, Matthias Rübsam, Carien M. Niessen, Leah C. Biggs, Sara A. Wickström

## Abstract

Formation of a bi-directional skin barrier is essential for organismal survival and maintenance of tissue homeostasis. Barrier formation requires positioning of functional tight junctions (TJ) to the most suprabasal viable layer of the epidermis through a mechanical circuit that is driven by generation of high tension at adherens junctions. However, what allows the sensing of tension build-up at these adhesions and how this tension is balanced to match the requirements of tissue mechanical properties is unclear. Here we show that the mechanosensitive ion channel Piezo1 is essential for the maturation of intercellular junctions into functional, continuous adhesions. Deletion of Piezo1 results in an imbalance of cell contractility and membrane tension, leading to a delay in adhesion maturation. Consequently, the requirement for Piezo1 activity can be bypassed by lowering contractility or elevating membrane tension. *In vivo*, Piezo1 function in adhesion integrity becomes essential only in aged mice where alterations in tissue mechanics lead to impaired TJs and barrier dysfunction. Collectively these studies reveal an essential function of Piezo1 in the timely establishment and maintenance of cell-cell junctions in the context of a mechanically tensed epidermis.

## Introduction

Epithelial cells line the surfaces of organs and generate a barrier to protect underlying tissues and maintain the physiological organ environment. Consequently, barrier dysfunction causes diseases ranging from infections to chronic inflammation and understanding the central mechanisms of barrier function is critical for the development of treatments to restore or maintain barriers ^1,2^. The skin epidermis is a multilayered stratified tissue that serves as the primary barrier between the body and the external environment. In mammalian skin the epidermis has two sets of physical barriers, the stratum corneum and tight junctions (TJs). The stratum corneum is the outermost epidermal layer, mainly consisting of terminally differentiated corneocytes and intercellular lipids. TJs are intercellular junctions present in the uppermost viable layer stratum granulosum II (SG2) ^3^. Together, these two barriers protect the body from environmental stresses such as irradiation, pathogens, chemical irritants, and mechanical stress, while also preventing water loss ^4–7^.

The formation of intercellular junctions is initiated by formation of so-called primordial adherens junctions that form an interdigitated “adhesion zipper” at the cell-cell interface. These zipper-like adhesions then evolve into spatially separated, continuous “belt-like” adherens junctions and TJs coinciding with epithelial polarization ^8,9^. TJs consist of claudin adhesion receptors, which polymerize into a network of intercellular strands creating the diffusion barrier ^10^. On the cytoplasmic side, a dense plaque of proteins is assembled, including zona occludens proteins (ZO) 1 and 2, that are already present in primordial adherens junctions and in TJs form a membrane-attached scaffold to facilitate formation and sub-apical positioning of claudin strands and to sequester cytoskeleton and signaling proteins ^11^. The tethering of claudins to actin through ZO-1 is dynamic and transient, likely facilitating cell shape adaptations during motility^12,13^. Importantly, the molecular mechanisms underlying the spatial assembly and formation of TJs from primordial adherens junctions involves an E-cadherin-driven mechanical signaling network that regulates the organization of the actomyosin cortex to stabilize TJs. Depletion of cadherins leads to decreased cortical tension and inability of cells to form TJs^14,15^. However, how cortical tension and contractility is sensed and balanced locally at cell-cell junctions is unclear.

Piezo1 is a mechanosensitive ion-channel located primarily at the plasma membrane and has been demonstrated to be activated by several mechanical stimuli such as stretching, shear stress, or by sensing changes in local curvature of the plasma membrane ^16–22^. Piezo1 has been shown to play a prominent role as a mechanosensor in development, homeostasis and disease ^23^. In the epithelium of *Xenopus* Piezo1-mediated calcium flashes have been implicated in maintaining an intact TJ barrier by regulating cortical actomyosin contraction ^24^. At the molecular level, Piezo1 has shown to localize at cell-matrix adhesions ^25^ and adherens junctions, where it has demonstrated to interact with E-cadherin, β-catenin, and filamentous (F)-actin ^26^. The role of Piezo1 in cell-cell junctions of the mammalian skin epidermis has not been studied.

Here we address the mechanisms by which cortical and membrane tension are balanced to facilitate formation and mechanical stability of intercellular junctions and the role of Piezo1 in this process. We show that timely transition of zipper-like primordial junctions into a mature, continuous, occludin-containing putative TJs requires Piezo1 mechanosensing. In the absence of Piezo1, intercellular junctions exhibit delayed maturation and decreased mechanical stability, which can be bypassed by modulating membrane or cortical tension. Consequently, *in vivo*, in the context of aging epidermis that exhibits elevated tissue stiffness, the absence of Piezo1 leads to the impairment of epidermal barrier function. Collectively these data demonstrate a role for Piezo1 in balancing membrane tension and cortical contraction, which is ultimately required to stabilize intercellular junctions and TJs to maintain skin barrier integrity.

## Results

### Intercellular junction maturation requires balancing cortex and membrane tension

To investigate the morphological changes at the cell-cell adhesion interface during intercellular adhesion maturation, we performed live imaging of mouse primary epidermal keratinocytes after switching cells to high Ca^2+^ **(Fig. 1a; Supplementary Movie 1).** Using ZO1-mEmerald as a marker, we quantified the transition from zipper-like primordial adherens junctions into continuous “belt-like”, junctions. We observed a temporally heterogeneous process with initiation of belt formation 80 min after adding Ca^2+^ into the growth medium, that was completed in all cells by 16 h post Ca^2+^ switch, concomitant with abrupt cellular flattening upon transition from zippers to belt-like adhesions **(Fig. 1a; Supplementary Movie 1).**

**Figure 1:**
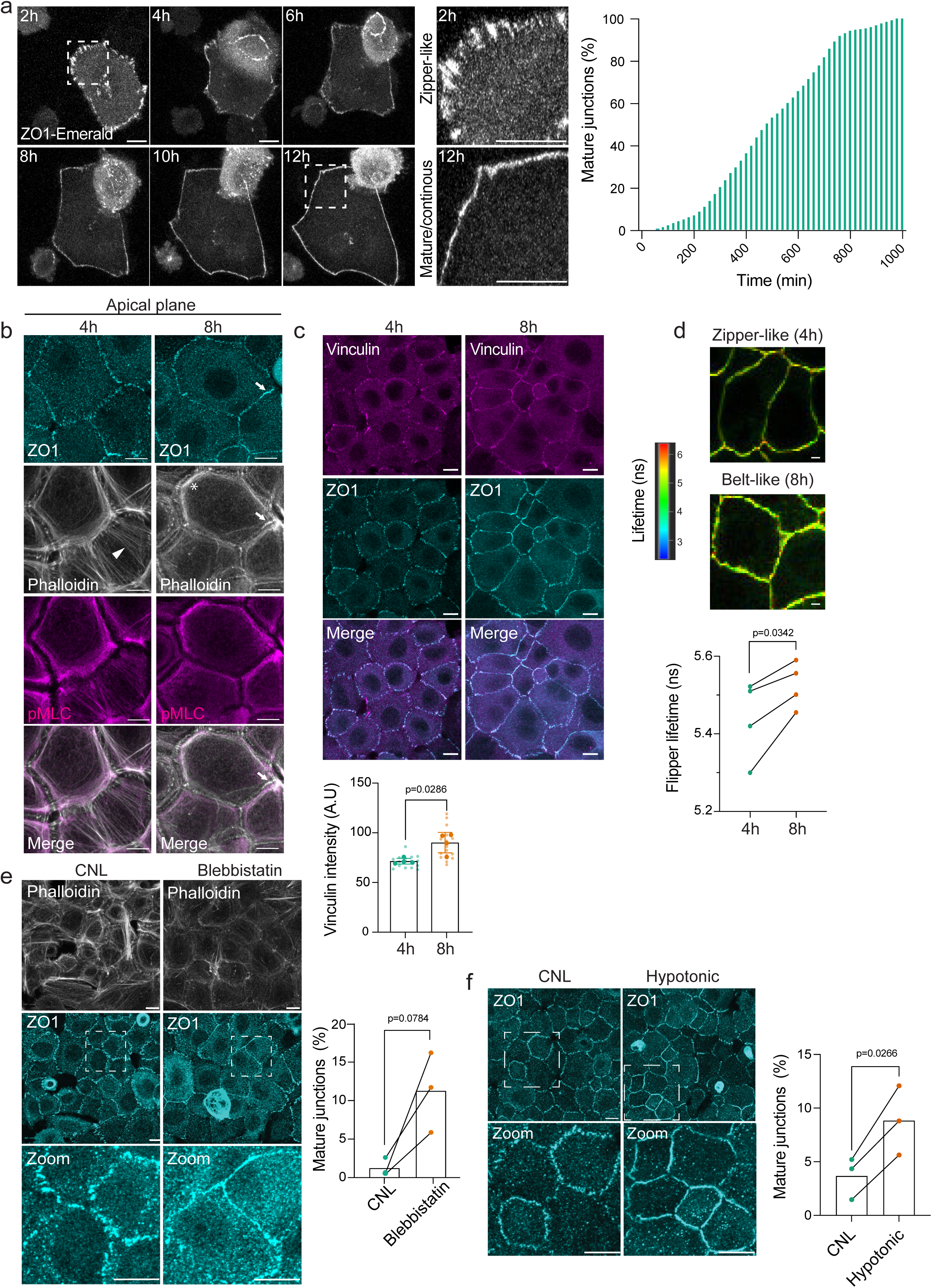
Intercellular junction maturation requires balancing membrane and cortical tension. **a)** Representative snapshots from live imaging data from ZO1-mEmerald expressing keratinocytes post Ca^2+^ switch at time points indicated. Dashed boxes indicate magnified areas (right panels). Quantification shows the proportion of cells with continuous junctions as a function of time (n=41 cells pooled across 3 independent experiments; scale bars 10 µm). **b)** Representative images of apical surface projections of ZO1, pMLC2 and phalloidin-stained primary keratinocytes fixed at 4 h and 8 h post Ca^2+^ switch. Note thin apical actin stress fibers (arrowhead) and punctate junctional actin (arrows) at 4 h and redistribution of actin into thick peripheral bundles at 8 h (asterisk) (n=3 independent experiments, scale bar 10 µm). **c)** Representative images of apical surface projections and quantification of vinculin (and ZO1-stained primary keratinocytes fixed at 4 h or 8 h after Ca^2+^ addition. Note increase in junctional vinculin intensity at 8 h (n=3 independent experiments; mean ± SD; Mann-Whitney, scale bar 10 µm). **d)** Representative images and quantification of fluorescence lifetime imaging of FLIPPER-TR. Note increase in FLIPPER-TR lifetime indicative of elevated membrane tension after 8 h Ca^2+^ (n=4 independent experiments with ≥ 50 membrane measurements per condition/experiment; paired t-test; scale bar 10 µm). **e)** Representative images of ZO1-stained cells treated with 5 µM blebbistatin. Dashed boxes indicate magnified areas (bottom panels). Quantification shows the faster maturation of junctions in blebbistatin treated cells (n =3 independent experiments; paired t-test, scale bar 10 µm). **f)** Representative images of ZO1-stained primary keratinocytes exposed to hypotonic medium (140 mOsm). Dashed boxes indicate magnified areas (bottom panels). Quantification shows accelerated maturation of cell-cell junctions in cells exposed to hypotonic medium (n =3 independent experiments; paired t-test, scale bar 10 µm).

Cell shape is mainly controlled by two opposing forces: cortical tension that reduces cell contacts^27^ and intercellular adhesion forces that increase the surface of contacts. We thus sought to analyze how these forces are balanced in space and time during adhesion maturation in keratinocytes. Adherens junctions can recruit cortical actin filaments directly, via α-catenin, or indirectly through proteins such as vinculin ^15,28–31^. Myosin activity and specifically non-muscle myosin IIA (NMIIA) then generates the mechanical tugging force necessary for cell-cell junction reinforcement and maintenance ^32^. At 4 h post Ca^2+^ application, apical F-actin was found organized in thin stress fibers and punctate junctional actin at zippers **(Fig. 1b)**. Concomitant with emergence of continuous junctions (8h), the stress fibers were replaced by thick actin bundles positioned perpendicularly to junctions **(Fig. 1b)**. In contrast to stress fibers and the perpendicular actin bundles, the actin at junctions was low in myosin activity (phosphorylated myosin light chain; pMLC2) **(Fig. 1b)**.

We next approximated stresses within the junctions themselves by quantifying their molecular links to actin - vinculin and α-catenin **(Fig. 1c)**. Quantification of vinculin intensity at the early zipper-like stage and mature belt-like stage indicated an increase in vinculin abundance **(Fig. 1c)**. Additional analysis of the tension-sensitive epitope (α18)-catenin ^33^ during the time course of maturation process confirmed the build-up of junctional stresses concomitant with adhesion maturation (**Supplementary Fig. 1a)**. This indicated that formation of belt-like adhesions was associated with initial contractility build-up by actomyosin stress fibers linked to junctions, followed by a switch to parallel actomyosin bundles and reduced contractility at adhesions, while the junctions themselves were stabilized in a stressed state indicated by a strengthened actin-junction link.

The results so far were consistent with previous observations that contact expansion from zippers to a belt requires collaborative regulation of adhesion tension and actomyosin cytoskeleton to lower interfacial tension at the contact^34,35^. While these models focus on cytoskeleton-based mechanisms, surface tension is also controlled by membrane tension^36^. Consistently, increased membrane tension has been shown to promote contacts between neighboring cell membranes to allow cadherin clustering^37^. Actomyosin-mediated cortex tension has further been shown to anticorrelate with membrane tension ^38,39^. We thus addressed changes in plasma membrane tension during junction maturation. To this end we performed fluorescence-lifetime imaging (FLIM) of a membrane tension reporter Flipper-TR ^40^, that we controlled by measuring lifetimes after hyper and hypotonic stress (**Supplementary Fig. S1b)**. FLIM imaging revealed an increase in Flipper-TR lifetime during the TJ maturation phase, indicative of increasing membrane tension **(Fig. 1d)**.

To test if balancing actomyosin and membrane tension are coordinating timely formation of continuous cell-cell junctions, we first inhibited myosin activity using blebbistatin or artificially elevated membrane tension by manipulating osmolarity during the time course of junction formation^41^. Indeed, treatment of keratinocytes with the myosin inhibitor blebbistatin ^42^ accelerated the transition of junctions from the immature zipper-like state to a continuous mature state **(Fig. 1e)**. Consistently, elevating membrane tension by inducing cell swelling with a mild hypotonic treatment led to accelerated maturation of cell-cell junctions **(Fig. 1f)**. Collectively, these data showed that actomyosin cortex tension and membrane tension play a critical role in coordinating the dynamics of cell-cell junctions transitioning from a zipper-like primordial state into a mature, belt-like configuration.

### Piezo1 regulates maturation of cell-cell junctions

To address the molecular mechanisms by which contractility and membrane tension are balanced, we hypothesized that this would require local sensing of membrane and cortex tension at the junction. A strong molecular candidate is the tension-sensitive ion channel Piezo1. To test the involvement of Piezo1, we generated an epidermis-specific Piezo1 knockout-mouse by crossing Piezo1 floxed/floxed mice ^43^ with a Keratin14-promoter-driven Cre^44^ line (from here on termed Piezo1-eKO). Primary epidermal keratinocytes from Piezo1-eKO and control mice were transfected with ZO1-mEmerald and live imaged. Quantification of cell-cell junction formation upon Ca^2+^ switch showed significantly delayed dynamics of junctional maturation in Piezo1-eKO cells **(Fig. 2a)**. While early zipper-like adhesion formation was comparable between Piezo1-eKO and control cells, Piezo1-eKO cells showed a substantial delay in transitioning into mature belt-like junctions **(Fig. 2a; Supplementary Movie 2)**. In addition, the Piezo1-eKO cells exhibited more dynamic cytoskeletal motion and irregularly shaped **(Fig. 2a; Supplementary Movie 2)**. Analysis of vinculin translocation to intercellular junctions showed reduced levels of vinculin at cell-cell contacts, but abundant vinculin at cell-matrix adhesions (**Supplementary Fig. S2a)**, indicating abnormal build-up of stresses at intercellular junctions of Piezo1-eKO cells.

**Figure 2.**
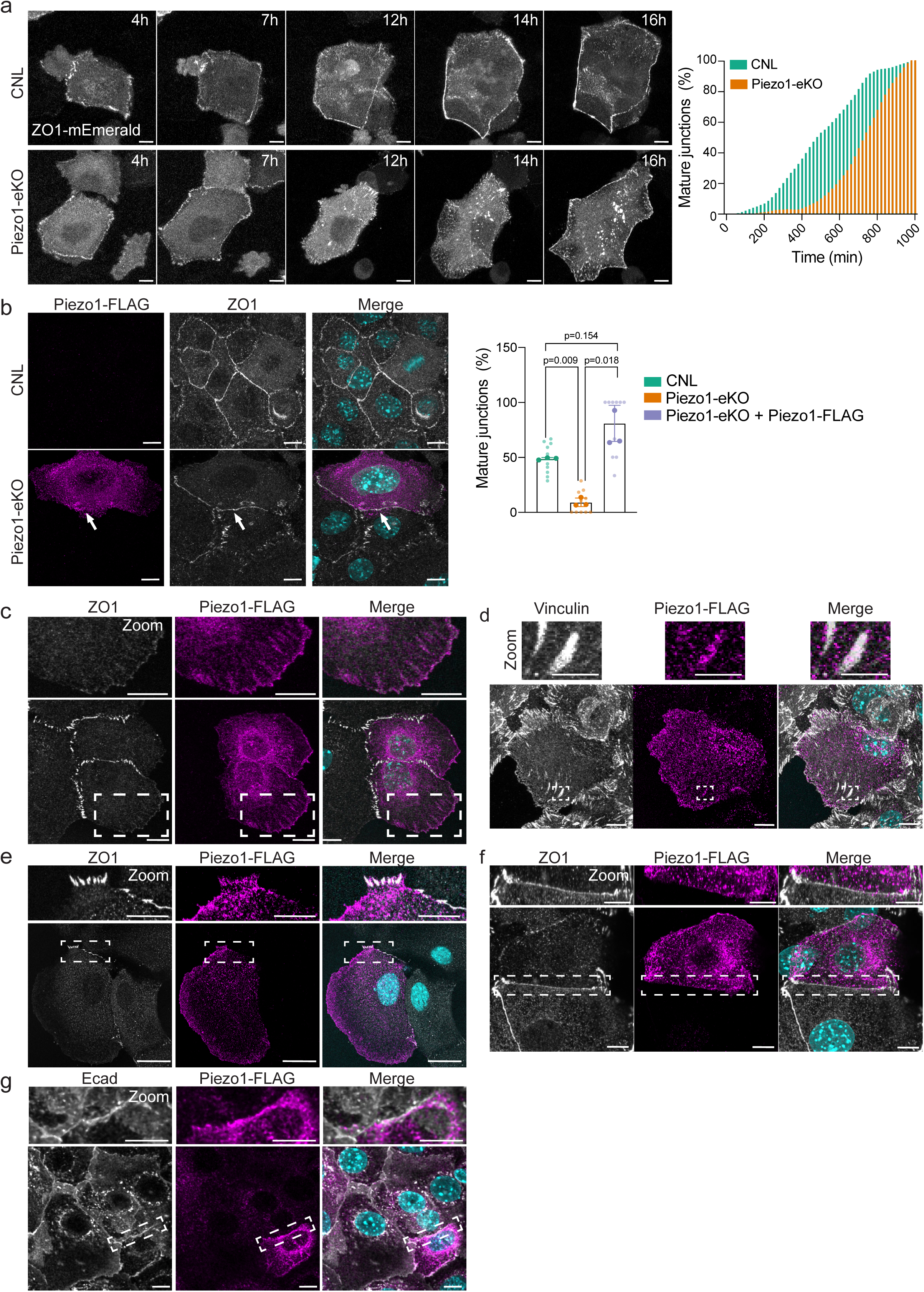
Piezo1 regulates maturation of cell-cell junctions. **a)** Representative live imaging snapshots of ZO1-mEmerald cell-cell junction assembly/maturation in CNL and Piezo1-eKO primary keratinocytes. Quantification shows relative proportion of cells with continuous junctions as a function of time. Note delay in junction maturation of Piezo1-eKO cells (n=41 (CNL)/ 30 (Piezo1-eKO) cells pooled across 3 independent experiments). **b)** Representative images of ZO1- and FLAG-stained CNL and Piezo1-eKO primary keratinocytes transfected with Piezo1-FLAG. Note that expression of Piezo1-FLAG in Piezo1-eKO rescues the dynamics of junction maturation (arrows) (n=3 independent experiments; one-way ANOVA; ns=not significant). **c)** Representative images of ZO1- and FLAG-stained Piezo1-eKO primary keratinocytes transfected with Piezo1-FLAG. Dashed boxes indicate magnified areas (top panels). Note localization of Piezo1-FLAG in focal adhesion-like structures. **d)** Representative images of vinculin- and FLAG-stained Piezo1-eKO primary keratinocytes transfected with Piezo1-FLAG. Dashed boxes indicate magnified areas (top panels). **e)** Representative images of ZO1- and FLAG-stained Piezo1-eKO primary keratinocytes transfected with Piezo1-FLAG. Note partial co-localization of ZO1 and Piezo1-FLAG in zipper-like adhesions. **f)** Representative images of ZO1- and FLAG-stained Piezo1-eKO primary keratinocytes transfected with Piezo1-FLAG. Note lack of co-localization of ZO-1 and Piezo1-FLAG at linear junctions. **g)** Representative images of E-cadherin- and FLAG-stained Piezo1-eKO primary keratinocytes transfected with Piezo1-FLAG. Dashed rectangles mark region of zoom-in. All scale bars 10 and 5 µm (zoom-in magnifications).

To confirm that this phenotype was not related to expression of exogenous, tagged ZO1, we performed additional immunofluorescence experiments at 3, 6, 12, and 48 h post Ca^2+^ switch where we stained cells for endogenous ZO1. Immunofluorescence analyses of fixed cells confirmed delayed maturation of Piezo1-eKO intercellular junctions 12 h post Ca^2+^ switch, and a delay in formation of occludin-positive TJ belts **(Supplementary Fig. S2b, c)**. Importantly, the delayed maturation phenotype caused by the lack of Piezo1 was rescued by reconstituting Piezo1 expression using Piezo1-FLAG, indicating that Piezo1 controls the maturation process **(Fig. 2b)**.

To understand if Piezo1 exerts its effect on intercellular junctions by directly localizing to these sites, we analyzed the distribution of Piezo1 during the time course of adhesion maturation. In the absence of well-functioning antibodies, we assayed the localization of the Piezo1-FLAG in the rescued Piezo1-eKO cells. Detection of Piezo1 with anti-FLAG antibody during the time course of cell-cell junction maturation revealed that prior to junction maturation, Piezo1 was enriched at focal adhesion-like structures together with vinculin **(Fig. 2c, d)**. Interestingly, Piezo1 also appeared to be partially co-distributed with ZO1 at zipper-like adhesions **(Fig. 2e)**. However, upon junction maturation, Piezo1 no longer localized to the belt-like junctions **(Fig. 2f, g).** Collectively, the data indicated that Piezo1 is required for cell junction maturation into junction belts and is localized to adhesion sites that are under high tension such as focal adhesions and zipper-like cell-cell junctions.

### Requirement of Piezo1 for cell-cell junction maturation can be bypassed by modulating membrane and cortical tension

Given that junction maturation requires balancing cortical and membrane tension, we next addressed whether Piezo1 regulates either of these parameters in order to promote timely cell-cell adhesion dynamics. FLIM imaging of the Flipper-TR membrane tension probe after 3 h of Ca^2+^ switch revealed reduced lifetimes in Piezo1-eKO cells compared to the control cells, indicative of lower membrane tension in the absence of Piezo1 **(Fig. 3a)**. Furthermore, we approximated cortex stiffness of Piezo1-eKO keratinocytes by using atomic force microscope-mediated force indentation spectroscopy (AFM) of a spherical bead cantilever to measure cortical stiffness^45,46^. Coinciding with the timescale of zipper-like adhesion formation at 3 h and 6 h post Ca^2+^ switch, which occurred normally also in Piezo1-eKO cells, loss of Piezo1 did not significantly affect the build-up of cortical stiffness at these time points **(Fig. 3b)**. However, at 12 h post Ca^2+^ switch when control cells had formed mature junction belts and cortical stiffness had plateaued, Piezo1-eKO keratinocytes showed a moderate increase in cortical stiffness **(Fig. 3b)**. Consistently, analyses of actomyosin contractility revealed persistence of actin stress fibers still at 8 hours post Ca^2+^ switch **(Fig. 3c),** suggesting a delay in balancing cortex tension.

**Figure 3.**
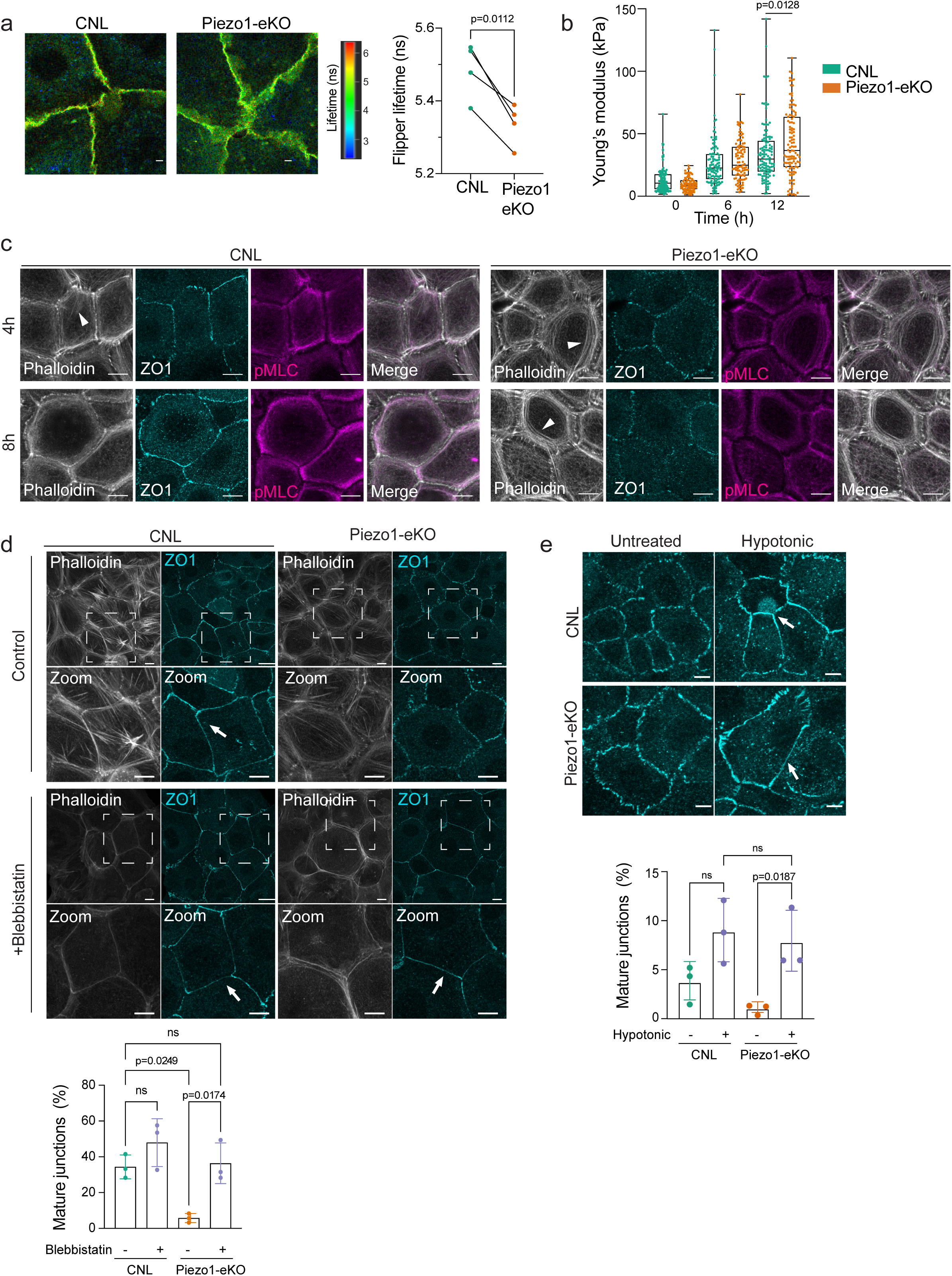
Requirement of Piezo1 for cell-cell junction maturation can be bypassed by modulating membrane and cortical tension. **a)** Representative images and quantification of fluorescence lifetime imaging of FLIPPER-TR from CNL and Piezo1-eKO primary keratinocytes. Note increase in FLIPPER-TR lifetime indicative of elevated membrane tension in Piezo1-eKO (n=4 independent experiments with ≥ 40 membrane measurements per condition/experiment; paired t-test; scale bar 10 µm). **b)** AFM force indentation spectroscopy of CNL and Piezo1-eKO primary keratinocytes before (0 h) and after Ca^2+^ switch. Note elevated elastic modulus in Piezo1-eKO at 12 h (n>88 force curves/condition pooled across three independent experiments; Kolmogorov-Smirnov). **c)** Representative images of apical surface projections of ZO1, pMLC2 and phalloidin-stained primary keratinocytes fixed at 4 h and 8 h post Ca^2+^ switch. Note delay in disassembling thin apical actin stress fibers (arrowheads) in Piezo1-eKO (n=3 independent experiments, scale bar 10 µm). **d)** Representative images and quantification of ZO1-stained CNL and Piezo1-eKO primary keratinocytes treated with 5µM blebbistatin for 6 h starting at 6 h of Ca^2+^ switch. Quantification shows faster maturation (arrows) of ZO1 based cell-cell junctions in blebbistatin-treated Piezo1-eKO cells (n=3 independent experiments; One-Way ANOVA/Tukey’s; ns=not significant, scale bar 10 µm). **e)** Representative images and quantification of ZO1-stained CNL and Piezo1-eKO primary keratinocytes treated with hypotonic media (140 mOsm). Quantification shows accelerated maturation of ZO1 based cell-cell junctions (arrows) in Piezo1-eKO cells in hypotonic conditions (n =3 independent experiments; One-way ANOVA/Tukey’s; ns=not significant, scale bar 10 µm).

To explore if the observed imbalance of actomyosin and membrane tension would explain the impaired junctional maturation in Piezo1-eKO cells, we proceeded to enhance membrane tension in these cells by increasing the osmotic pressure, and to reduce cortical contractility using blebbistatin. Strikingly, reducing cortical contractility rescued the junctional maturation defect **(Fig. 3d)**. Further, increasing membrane tension by hypotonic shock also facilitated timely maturation of cell-cell junctions in Piezo1-eKO cells **(Fig. 3e)**. Collectively these experiments showed that Piezo1 is required to balance membrane and cortex tension to facilitate junction maturation into belts.

### Aging leads to defects in tight junction maturation and barrier function of Piezo1-eKO mice

To understand if the observed role of Piezo1 in junction dynamics was relevant for epidermal development and homeostasis *in vivo*, we first examined Piezo1 expression within the stratified epidermal layers by combining RNA *in situ* hybridization (RNAscope) with immunofluorescence. Staining of wild type back skin sections with markers for basal stem cells (Keratin-14) and suprabasal differentiated cells (Keratin-10) and Piezo1 RNA revealed Piezo1 expression in both layers, with slightly more abundant Piezo1 mRNA in the suprabasal layers (**Supplementary Fig. S3a**). Consistent with the RNAscope results, re-analyses of published single cell RNA sequencing data from adult back skin^47^ showed highest expression of Piezo1 in the differentiated layers of the epidermis **(Supplementary Fig. S3b)**. Further, triggering adhesion maturation and terminal differentiation *in vitro* using Ca^2+^ switch also led to the increased mRNA levels of Piezo1 **(Supplementary Fig. S3c, qPCR)**, indicating that Piezo1 expression is temporally correlated with keratinocyte differentiation and/or junction maturation.

Next, we addressed the role of Piezo1 in regulation of epidermal homeostasis and intercellular junction formation *in vivo* by analyzing Piezo1-eKO skin. Piezo1-deficient skin showed no gross skin or hair phenotype, and histological analysis of young adult back skin mice (2-6 months old) revealed no morphological differences (**Supplementary Fig. S3d**). For optimal visualization of cell-cell junctions, we adapted used epidermal whole-mounts^15^. Staining of young adult ear whole mounts revealed no significant differences in TJ morphology as marked by ZO1 and occludin stainings or in overall cell morphologies (**Supplementary Fig. S3e**).

Given the role of Piezo1 in mechanosensing and the increase in tissue stiffness of aged mouse skin ^48^, we proceeded to analyze the changes to the epidermal phenotype during aging. Indeed, at 1 year of age, ear whole mounts showed reduced junctional ZO1 intensity within the SG2 layer **(Fig. 4a)**. We also observed reduced levels of vinculin at cell-cell contacts in the SG2 layer of Piezo1-eKO skin **(Fig. 4a)**, similar to the phenotype observed in cell culture. Quantification of SG2 cell shape and TJ morphology revealed more irregular cell shapes as well as less straight junctions in aged Piezo1-eKO mice **(Fig. 4b, c)**.

**Figure 4.**
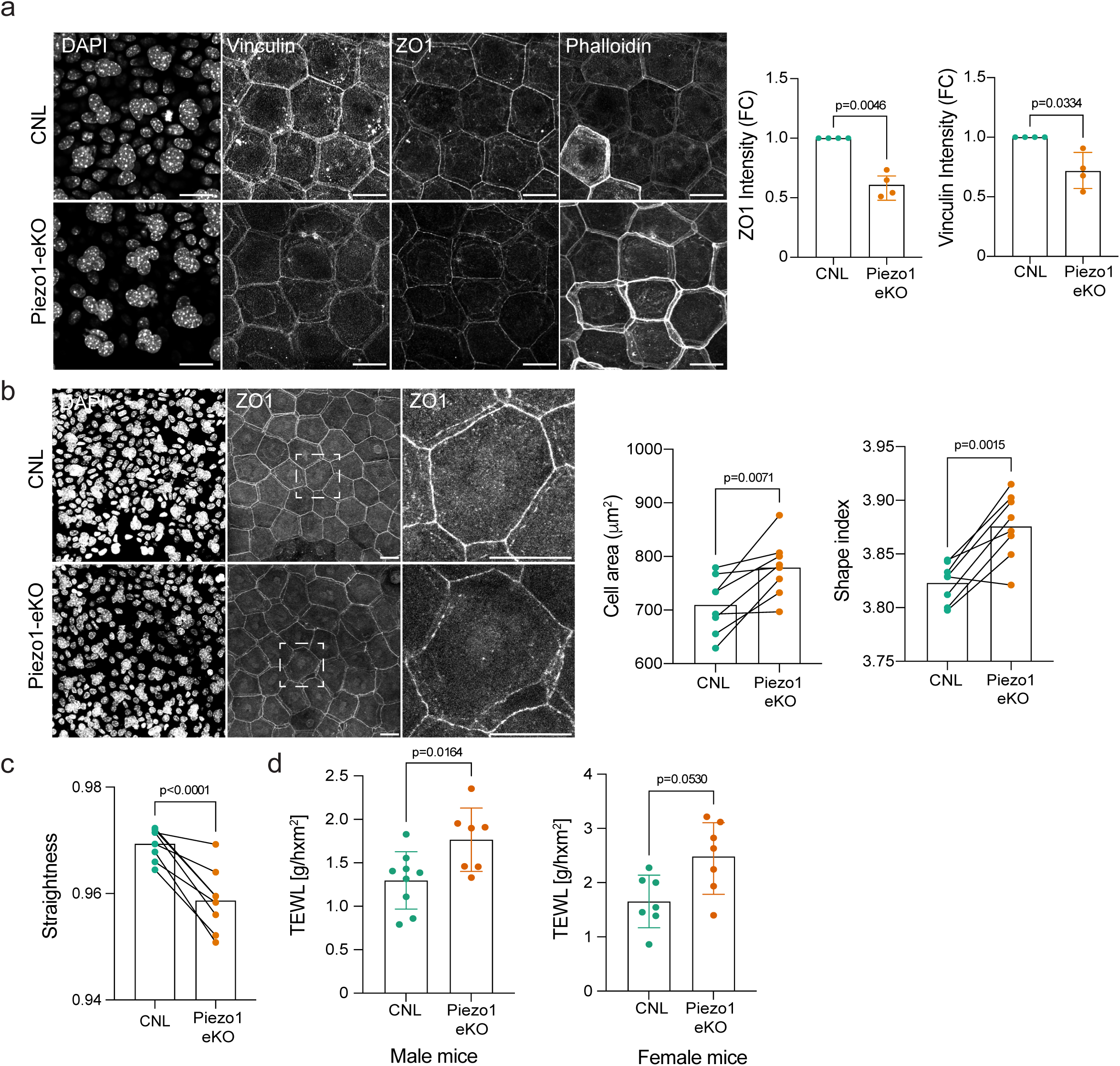
Aging leads to defects in tight junction maturation and barrier function of Piezo1-eKO mice. **a)** Representative images and quantification of vinculin, ZO1 and Phalloidin-stained ear whole mounts from 1 y-old CNL and Piezo1-eKO mice. Note decrease in junctional vinculin and ZO1 intensity in Piezo1-eKO (n=4 mice/genotype; one-sample t-test; scale bar 20 µm). **b)** Representative images and quantification of cell shape from ZO1-stained ear whole mount images of CNL and Piezo1-eKO <1-year old mice stained for ZO1. Dashed rectangle indicates zoomed-in area (n=8-10 mice/genotype; paired t-test, scale bars 20µm). **c)** Quantifications of junction straightness (n=8-10 mice/genotype; paired t-test). **d)** Quantifications of transepidermal water loss measurements from CNL and Piezo1-eKO 1-y old male and female mice back skin (n=9 (CNL male) / 7 mice (other groups); Mann-Whitney).

Given the critical role of TJs in establishing the bi-directional permeability barrier, we proceeded to test the functionality of these abnormal TJs by measuring transepidermal water loss (TEWL) in newborn, young adult, and 1 year old mice. Consistent with the morphologically unaltered TJs, no barrier impairment was observed in the early postnatal stage P1-3 **(Supplementary Fig. S3f)** or young adult mice **(Supplementary Fig. S3g, h)**. In contrast, and consistent with the abnormal TJs and cell shapes, male and female 1year old Piezo1-eKO mice displayed an increase in transepidermal water-loss, indicating that the barrier function in Piezo1-eKO mice is compromised **(Fig. 4d)**. Collectively, these findings indicate that Piezo1 is required for efficient TJ maturation and barrier function in aged mice, possibly due to increased tissue stiffness.

### Deletion of Piezo1 results in impaired epidermal homeostasis and tissue fragility

To understand if the observed barrier dysfunction was consequential for skin homeostasis in aged mice, we performed H/E staining of Piezo1-eKO aged mice. Consistent with the reduced barrier function in aged mice, H/E-stained epidermal sections of aged Piezo1-eKO mice exhibited epidermal hyperthickening **(Fig. 5a).** The hyperthickening was associated with hyperproliferation of the basal cells within the epidermis, whereas no hyperproliferation was detected in young mice. **(Fig. 5b, Supplementary Fig. S4a)**. As the hyperthickening and hyperproliferation was only observed in aged basal cells, we hypothesized that the observed abnormal proliferation was secondary to the barrier function rather than a cell autonomous phenotype. In support of this notion, cultured Piezo1-eKO keratinocytes did not display proliferation abnormalities **(Supplementary Fig. S4b)**, indicative of a cell non-autonomous effect.

**Figure 5.**
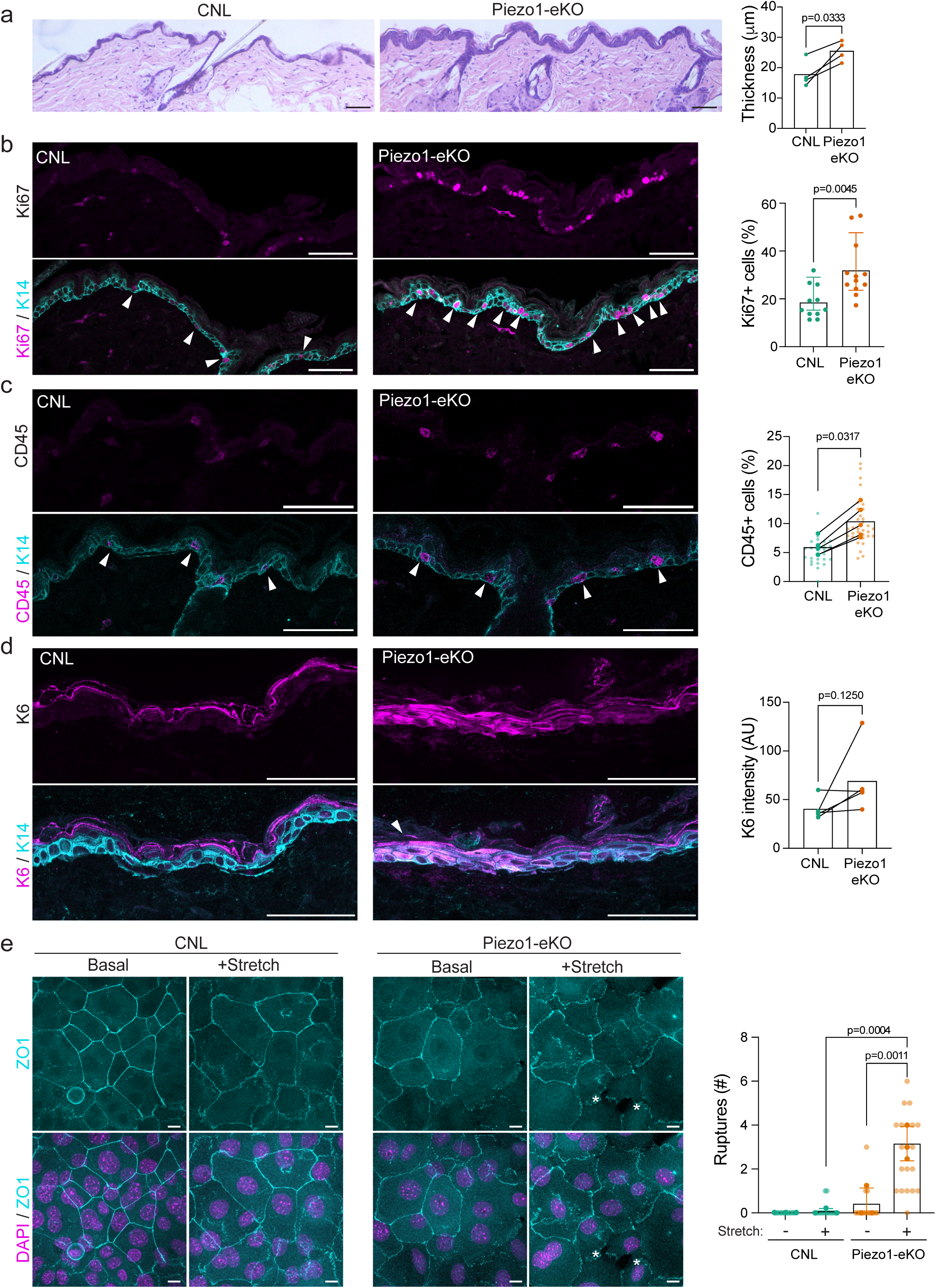
Deletion of Piezo1 results in impaired epidermal homeostasis and tissue fragility. **a)** Representative images of H/E-stained back skin sections and quantification of epidermal thickness from 1 y-old CNL and Piezo1-eKO back skin (n= 4 mice/genotype; unpaired t-test, scale bar 100 µm). **b)** Representative images and quantification of 1 y-old CNL and Piezo1-eKO mouse back skin stained for Ki-67 (magenta) and Keratin-14 (cyan). Note increase in Ki-67 positive cells (arrowheads) in epidermis of Piezo1-eKO mice (n=11 (CNL) and 12 (Piezo1-eKO); mean± SD; Mann-Whitney, scale bar 50 µm). **c)** Representative images and quantification of 1 y-old CNL and Piezo1eKO back skin stained for CD45 (magenta) and Keratin-14 (K14) (cyan). Note increased number CD45 positive cells (arrowheads) in Piezo1-eKO mice (n=5 mice/genotype; Mann-Whitney; scale bar 50 µm). **d)** Representative images and quantification of 1 y-old CNL and Piezo1-eKO back skin stained for Keratin-6 (K6) (magenta) and Keratin-14 (cyan). Note increased expression of Keratin 6 (arrowheads) in epidermis of Piezo1-eKO mice (n=5 mice/genotype; Mann-Whitney, scale bar 50 µm). **e)** Representative images and quantification of ZO1-stained biaxially stretched CNL and Piezo1-eKO primary keratinocyte monolayers. Note fractures in Piezo1-eKO monolayers (asterisks) upon stretch (n =3 independent experiments; One-way ANOVA/Tukey’s; scale bar 10 µm).

An impairment in skin barrier may trigger an inflammatory response in resident immune cells ^49^. To investigate if this is the case, we immunostained skin sections of aged Piezo1-eKO mice with leukocyte and macrophage markers, CD45 and F4/80, respectively. As anticipated, the absence of Piezo1 in the epidermis led to increased levels of resident leukocytes and macrophages **(Fig. 5c, Supplementary Fig S4c)**. As further indication of barrier dysfunction, we found that Piezo1-eKO mice exhibited elevated levels of epidermal stress as quantified by increased levels of the stress-associated Keratin-6, normally not expressed in healthy epidermis ^50^ **(Fig. 5d)**. Together, these results indicate that Piezo1-eKO mice exhibit a mild age-dependent barrier defect accompanied by hyperproliferation, inflammation and stress in the epidermis.

Finally, we asked why the TJ phenotype only manifests in aged mice. We hypothesized that aging-induced tissue-stiffening could increase the demands for mechanosensing to balance cortical contractility and membrane tension to efficiently form TJ ^48,51^ **(Fig 5 a-d)**. To test this, we experimentally elevated cortical contractility *in vitro* in primary keratinocytes by performing biaxial cell monolayer stretching. Analyses of junctional integrity upon application of mechanical stress revealed that whereas the cell-cell junctions of control monolayers were largely unaffected, Piezo1-eKO monolayers showed more ruptures at junctions, resulting in gaps in the monolayer **(Fig. 5e).** This indicates that Piezo1 is required for mechanical stability of cell-cell junctions, possibly explaining the appearance of the barrier defect only in the aged, mechanically stiff tissue.

## Discussion

Local changes in the balance of forces between cells, associated with events such as changes in tissue mechanics, are central to epithelial homeostasis^52,53^. Our current data suggest that balancing between cortical and membrane tension is required for junction maturation from zipper-like primordial junctions into mature TJ belts. We find that this balancing is mediated by the mechanosensitive ion channel Piezo1 in skin keratinocytes. While formation of primordial junctions occurs normally in the absence of Piezo1, Piezo1-deficients cells exhibit a strong delay in transitioning these actomyosin-linked zipper-like adhesions into mature adhesion belts. This delay can be bypassed by decreasing actomyosin cortex tension or increasing plasma membrane tension. Interestingly, the adhesion maturation defect becomes relevant for *in vivo* epidermal homeostasis only in aged mice, where an impaired TJ barrier is associated with epidermal hyperproliferation and hyperthickening, epidermal stress and an inflammatory response.

The switch from zipper- to belt-like intercellular junctions is tightly coupled to dramatic reorganization of the actomyosin cytoskeleton. In zippers, actomyosin stress fibers perpendicularly associate with adhesions to generate tension at junctions. Upon formation of belts these radially oriented stress fibers transition into peri-junctional actin bundles^54,55^. Thus, profound reorganization of actomyosin is crucial for this transition, and it is known that this switch is regulated both by local RhoA and Cdc42 activation downstream of the polarity master regulator aPKC^56,57^. However, what defines the precise timing of activation of these pathways has remained unclear. We propose that Piezo1, which transiently localizes to areas of high tension, focal adhesions and zipper-like adherens junctions, acts as a sensor for local tension build-up to determine the timing of the transition. Consequently, Piezo-deficient cells display prolongation of the zipper-like state. In this respect it is interesting to note that Piezo channels have been implicated in RhoA activity regulation in other mechanosensory processes such as focal adhesion maturation in mammalian cell lines or TJ repair in Drosophila, providing a potential downstream mechanism for this effect^58–60^.

While the role of dynamic actomyosin regulation in junction maturation is quite well understood, the role of membrane tension has been elusive. Membrane tension is defined as the force needed for the in-plane deformation of a unit length of membrane, and is driven by shifts in the equilibrium distance between phospholipids in the lipid bilayer^61^. The large size and curved architecture of Piezo1 trimers has been shown to render Piezo1 directly sensitive to lateral plasma membrane tension^62–65^. In addition to membrane tension, several studies indicate that the cytoskeleton gates activation of Piezo channels, albeit the direct link between Piezo1 and cytoskeleton is under debate^66–71^. This dual sensitivity places Piezo1 in an ideal position to integrate membrane tension and cortical tension sensitivity. Interestingly, we observe that Piezo1 is partially located at E-cadherin adherence junctions as well as focal adhesions, but is absent from belt-like junctions. This is consistent with previous work showing Piezo1 preferential localization at regions of high actomyosin tension: the contractile rear of migrating keratinocytes to controlling the speed of wound healing^72^, within focal adhesions of various cell types^59,73^, and to the intercellular bridge during cytokinesis^74^. This leads us to postulate that Piezo1 is localized at areas of high cortex tension, and it translocates away from the belt-like adhesion as the cortical contractility becomes lower.

Notably, the cortical cytoskeleton can control the length scale of membrane tension propagation, thereby indirectly controlling Piezo1 activation via the lipid bilayer^75–79^. Interestingly, while sensing membrane tension, Piezo1 has been shown to be involved in a feedback loop where Piezo1 activation leads to elevated membrane tension^80^. Finally, it has been shown during a sustained stimulus, Piezo1 currents will decay, preventing them from responding to an additional mechanical stimulus or steady state membrane tension^81^. These properties could collectively explain our observations that Piezo1 is required from the rapid, well-timed transition of high actomyosin tension generated zipper like junctions by localizing transiently specifically at these sites to sense local membrane tension. Once activated, downstream RhoA or Cdc42 signalling would then catalyze the necessary actomyosin rearrangement to generate the mature belt structures, further facilitated by increased membrane tension to bring the plasma membranes together. The reported decay mechanism of Piezo1 activity would subsequently inactivate the channels and the decrease in local actomyosin would facilitate Piezo1 localization away from these structures. Rigorously testing this model is an important task for future work.

Surprisingly, while the phenotype of Piezo1 junction maturation is a robust phenomenon in primary keratinocytes *in vitro*, observed consistently regardless of the age of the mouse from which the cells are isolated from, *in vivo* the phenotype only becomes obvious in aged mice. Tissue culture plastic is orders of magnitudes stiffer than *in vivo* tissues^82^, including skin, and previous studies have shown that the epidermis becomes stiffer with age^48,83^. This would imply that the importance of Piezo1 in local mechanosensing at the junctions becomes essential only at high levels of tissue tension and associated actomyosin contractility. Consistently, reducing contractility *in vitro* facilities timely adhesion maturation in the absence of Piezo1, whereas increasing monolayer tension by stretch makes adhesions less stable.

Collectively these data implicate Piezo1 in sensing membrane tension at sites of high actomyosin contractility, thereby integrating these two processes to regulate timely adhesion maturation, particularly in tissues that are under high mechanical tension.

## Materials and Methods

### Mouse strains

All mouse studies were approved and carried out in accordance with the guidelines of the Finnish national animal experimentation board (ELLA) under permit number ESAVI/22531/2018. C57BL/6J mice with homozygous floxed Piezo1 alleles (Piezo1^fl/fl^, Jackson Labs stock no. 029213), were bred with K14-Cre mice ^84^ to achieve Piezo1^fl/fl^-K14-Cre mice with epidermis-specific deletion. No gender differences were detected in phenotypes with the exception of TEWL, so both males and females were analyzed. Piezo1^fl/fl^ and Piezo1^fl/wt^ (collectively termed as control (CNL)) sex-matched littermates were used as control.

### Isolation and culture of primary keratinocytes

Epidermal keratinocytes were isolated from adult (3 months old) control and Piezo1-eKO mice. Following skin isolation, epidermal single cell suspensions were generated by incubating skin pieces in 0.8% trypsin (Gibco) for 45-50 min. Following trypsinization, the epidermis was separated from dermis by scraping with small forceps. The epidermal-cell suspension was pipetted up and down several times and filtered through a 70 µm cell strainer. Cells were resuspended in Keratinocyte Growth Medium (MEM Spinner’s modification (Sigma), 5 µg ml^−1^ insulin (Sigma), 10 ng ml^−1^ EGF (Sigma), 10 µg ml^−1^ transferrin (Sigma), 10 µM phosphoethanolamine (Sigma), 10 µM ethanolamine (Sigma), 0.36 µg ml^−1^ hydrocortisone (Calbiochem), 2 mM glutamine (Gibco), 100 U ml^−1^ penicillin, 100 µg ml^−1^ streptomycin (Gibco), 10% chelated fetal calf serum (Gibco), 5 µM Y27632, 20 ng ml^−1^ mouse recombinant vascular endothelial growth factor and 20 ng ml^−1^ human recombinant fibroblast growth factor-2 (all from Miltenyi Biotec) as described ^85^ and plated on pre-coated (Collagen I (10 µg ml^−1^ Millipore), Fibronectin (100 µg ml^−1^ Millipore) culture dishes.

### Transfection and drug treatments

hPiezo1-3xFLAG was from Genecopeia (EXZ6777-Lv181). ZO1-mEmerald was from Addgene (# 54316). For transfections, cultured keratinocytes were trypsinized using 0.5% trypsin (Gibco 15090-046) with 0.5mM EDTA. 300 000 cells were seeded in 8-well chamber slides (Lab-Tek) and cultured in KGM for 1-2 days until 80-90% confluent. Transfections were performed with Lipofectamine3000 according to the manufacturer’s protocol. After 6-8h transfection, the media was switched to Defined Keratinocyte Serum Free Medium (KSFM) (Gibco, Cat. #107850-12) supplemented with 200 µM Ca^2+^ to induce the formation of junctions.

For hypotonic treatment, the KSFM medium was diluted with sterilized miliQ water in 1:1 ratio leading to approximate molarity of 130-160 mOsm. Ca^2+^ was added to a 200 µM final concentration. For blebbistatin treatment, primary keratinocytes were first cultured in KSFM+200 µM Ca^2+^ for 6 h to induce the initial formation of immature junctions. Subsequently, 5 µM blebbistatin (Sigma B0560) was added in the culture media and cells were cultured for another 6 hours.

### Immunofluorescence

Cells were cultured on glass coverslips and fixed in 4% paraformaldehyde (PFA) in phosphate buffered saline (PBS) for 10 min at 37°C. After rinsing, cells were incubated with blocking solution (3% BSA and 5% normal goat serum (NGS), 0.3% Triton-X in PBS) for 30-45 min at room temperature. After blocking, cells were incubated overnight at 4°C with primary antibodies diluted in 1% BSA, 0.3% Triton-X in PBS. After multiple washes in PBS, secondary antibodies were diluted in 1% BSA, 0.3% Triton-X in PBS and incubated for 30 min at room temperature. After repeated washes, coverslips were mounted on objective slides with Elvanol mounting medium.

Tissue biopsies were fixed in 4% PFA, dehydrated through graded ethanol series, and embedded in paraffin using standard protocols and sectioned. Sections were deparaffinized using a graded alcohol series, blocked in 3% BSA and 5% NGS, after which antigens were retrieved using DAKO antigen retrieval solution (pH6 and pH9). The sections were incubated with primary antibodies diluted in Dako Antibody Diluent overnight at 4°C. After multiple washes in PBS, secondary antibodies were diluted in % BSA, 0.3% Triton-X in PBS and incubated for 30 min at room temperature. After washing, slides were mounted in Elvanol.

The following primary antibodies were used in this study: rabbit polyclonal anti-ZO-1 (Invitrogen 40-2200; 1:250), rat monoclonal anti-tension sensitive α18 epitope of α-catenin ^33^ (kind gift from Akira Nagafuchi; 1:3000,), rabbit polyclonal anti-Keratin-6 (Covance #PRB-169P; 1:500), mouse monoclonal anti-Vinculin (Millipore MAB3574; 1:250), guineapig polyclonal anti-Keratin K14 (Progen GP-CK14; 1:500), rat monoclonal anti-F4/80 (BMA Biomedicals T-2028; 1:200), mouse monoclonal anti-CD45 (eBioscience 30-F11; 1:200), mouse monoclonal anti-Ki67 (Cell Signaling #9449; 1:400), mouse monoclonal anti-E-cadherin (BD Biosciences #610181; 1:300);

The secondary antibodies were: AlexaFluor 488, 594, or 647 (Invitrogen; 1:500). Alexa Fluor 647 Phalloidin (Invitrogen A22287; 1:400) was used to stain F-actin. Nuclei were detected by 4’,6-diamidino-2-phenylindole (DAPI).

### Preparation of epidermal ear whole mounts

Epidermal ear whole mounts were prepared from young and old mice as described previously ^15^. Briefly, ears were dissected into dorsal and ventral pieces, cartilage was removed, and the inner (ventral) side of the ear was floated on a 5 µM Dispase II (Sigma D4693) solution at 37°C for 10-15 min. The epidermal layer was peeled off and fixed on ice with 4% PFA for 10 min, rinsed in PBS and permeabilized with 0.5% TritonX100/PBS for 1h at room temperature. Following permeabilization, the epidermal sheet was blocked with 3% BSA and 5% NGS after which primary antibodies were diluted in blocking solution and incubated overnight at 4°C. Secondary antibodies, DAPI and Phalloidin were subsequently incubated for 45min at room temperature, after which tissues were rinsed extensively and mounted on objective slides using Elvanol.

### Microscopy

Confocal images were obtained using Leica TCS SP8X with white light laser and Leica Stellaris 8 FALCON/DLS with 20X 0.75 NA, 40x 1.30 NA, and 63X 1.40 NA objectives. Live imaging was performed using LSM980 (Zeiss) confocal microscope equipped with an Airyscan2 detector, and an environmental chamber set at 37 °C, 5% CO using a 40X immersion objective. Images were acquired with Zeiss ZEN (Zeiss ZEN version 3.5) software where ‘joint deconvolution’ processing was performed post image acquisition.

### Image processing and analysis

The transition from zipper-like adhesions to mature continuous intercellular junctions were quantified manually. For quantifying intensities at junctions, max projection images were generated, and region of interests (ROIs) were restricted to ZO1-positive junctions. For cell shape, ROIs were generated for cell outlines, the area and shape were extracted, and cell shape was calculated by shape index (Perimeter / ✓Area). For quantifying junctional length and straightness, ROIs were drawn over ZO1 junctions manually, length and ferret length values were extracted, and junctional straightness was calculated from ferret length / length. Vinculin apical to basal ratio was calculated by dividing vinculin mean intensity from junctional focal plane with basal focal plane.

Quantification of Ki-67-, F4/80- and CD45-positive cells were counted manually. For quantification of Keratin.6 intensity in the epidermis, ROIs were generated for the epidermal area and intensities were extracted within these ROIs.

### Transepidermal water loss measurement

Transepidermal water loss was assessed as described by using noninvasive probe Tewameter TM300 (Courage+Khazaka, Cologne, Germany) ^86,87^, according to the manufacturer’s instructions. The dorsal skin hairs of mice were shaved prior to measure TEWL. Mice were anesthetized using isoflurane during TEWL measurements. TEWL measurements were averaged at 1-second intervals for a 1–2-minute period. For consistency, TEWL was measured by placing the probe at the same location on the dorsal skin of the mouse each time and the same person made all TEWL measurements.

### Histological stainings

For histological sections staining, Hematoxylin and Eosin Stain kit (Nordic BioSite Cat. # KSC-1ZN4SF-1) was used according to the manufacturer’s protocol. After staining, the sections were mounted in Entellan.

### RNA in situ hybridization combined with immunofluorescence

RNA in situ hybridization was performed using RNAscope technology following manufactured instructions (ACD, Multiplex Fluorescent v2 Assay combined with Immunofluorescence - Integrated Co-Detection Workflow). Briefly, tissue sections (5 μm thickness) were deparaffinized in xylene, followed by dehydration in 100% ethanol and incubated in hydrogen peroxide for 10 min at RT and placed in RNAscope Co-Detection Antigen Retrieval solution (ACD; #323165) for 15 min at 100 °C in a pressure cooker. Primary antibodies were incubated overnight at 4°C in a humidified chamber. Post fixation in 4% PFA was performed for 30 min at room temperature, after which RNAscope Protease Plus (ACD; #322381); was added to each sample and incubated at 40° C for 30 min. The Piezo1-RNAscope probe (ACD; #500511-C2) was hybridized following manufacturer instructions using RNAscope Multiplex fluorescent v2 assay (ACD; #323110) followed by detection with TSA plus Fluorescein (Akoya) fluorophore. All the steps were performed in an Hyb-EZ II oven (ACD). After hybridization, the samples were incubated with secondary antibodies for 30min at room temperature (AlexaFluor 594, or 647 (Invitrogen; 1:500). The samples were mounted in Elvanol and imaged using a Zeiss LSM 980 confocal, with Zeiss ZEN (Zeiss ZEN version 3.5) software with a 40X water immersion objective.

Piezo1 RNAscope images were quantified by drawing region of interest mask based on Keratin-14 positive or Keratin-10-positive cells after fluorescent foci of Piezo1 mRNA were quantified using the Spot Counter tool in Fiji.

### Force indentation spectroscopy (AFM)

AFM measurements were performed on cell monolayers plated glass bottom dishes using JPK NanoWizard 2 (Bruker Nano) atomic force microscope mounted on a Nikon Eclipse Ti inverted microscope and operated with JPK SPM Control Software v5. Spherical Silicon Nitride 5µm cantilevers (MLCT, Bruker Daltonics) with a nominal spring constant of 0.065 Nm^−1^ were used for the nanoindentation experiments of the apical surface/cortex of cells. For all indentation experiments, forces of up to 8.946 nN were applied, and the velocities of cantilever approach and retraction were kept constant at 10 μm s^−1^. All analyses were performed with JPK Data Processing Software (Bruker Nano). Prior to fitting the Hertz model corrected by the tip geometry to obtain Young’s Modulus (Poisson’s ratio of 0.5), the offset was removed from the baseline, contact point was identified, and cantilever bending was subtracted from all force curves.

### Membrane tension measurements

Flipper-TR fluorescent tension probe (Spirochrome, SC020) was utilized to quantify membrane tension ^40^. Cells were cultured on glass bottom dishes and switched to KSFM + 200 μM Ca^2+^ for 3 h, 4h or 8h. Flipper-TR (1 μM) was applied 10-15 minutes prior to measurements. FLIM imaging was performed using a Leica Stellaris confocal microscope equipped with a FLIM module. Leica FALCON/FLIM software was used to record the data in photon-counting mode using a HC PL APO CS2 63x 1.40 NA oil immersion objective. Excitation was performed using a pulsed 488 nm laser operating at 20 MHz with emission collected through bandpass 575/625 nm with a HyD X2 detector. Leica FLIM software was used to fit fluorescence decay data from regions of interest to a two-exponential model. The longest lifetime with the higher fit amplitude was used to quantify membrane tension.

### Mechanical stretching

For biaxial stretch, culture plates with a silicon elastomer membrane (Bioflex; FlexCell International Corporation) were coated with fibronectin (10 μg/ml) in PBS at 37°C overnight prior to cell seeding. 1.5 M cells per elastomer were seeded 24-48 h prior to experiment start to obtain 100% confluency, and 12 h before initiation of stretch culture medium was replaced by KSFM+200 μM Ca^2+^. Cells were then exposed to cyclic mechanical strain using the Flexcell Tension System (FX4000; FlexCell International Corporation) at 20 % elongation, 0.6 Hz frequency for 30min. After stress cessation, the cells were immediately fixed in culture medium using 8% PFA and immunostained as described above.

### qPCR

RNA was isolated using the Nucleospin RNA Plus kit (Macherey&Nagel), after which cDNA was synthesized using the iScript cDNA synthesis kit (Bio Rad). qPCR was performed on the StepOne Plus Real Time PCR System (Applied Biosystems) using the PowerUp SYBR Green Master Mix (Applied biosystems). Gene expression changes were calculated following normalization to GAPDH using the comparative Ct (cycle threshold) method.

The primers used are provided in the following table:

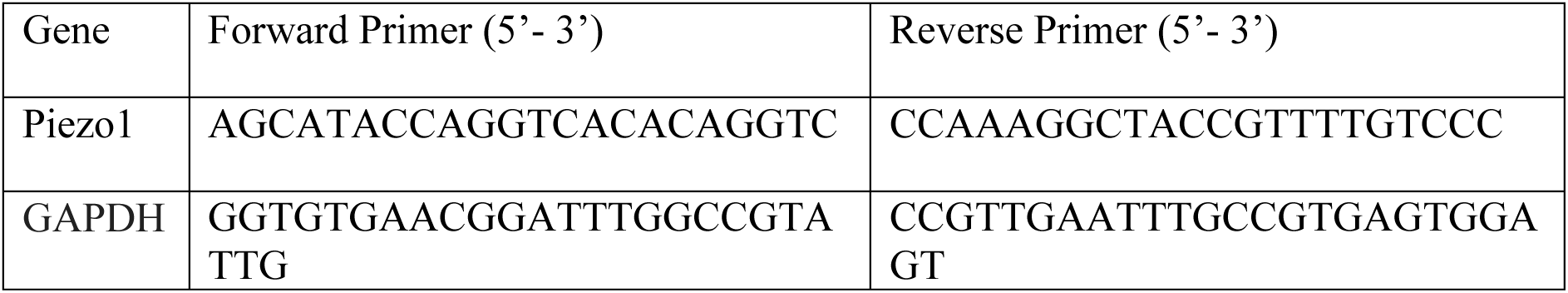

### Statistics and reproducibility

Statistical analyses were performed using GraphPad Prism software (GraphPad, version 10). Statistical significance was determined by the specific tests indicated in the corresponding figure legends. In all cases where a test for normally distributed data was used, normal distribution was confirmed with the Kolmogorov–Smirnov test (α = 0.05). All experiments presented in the manuscript were repeated at least in 3 independent biological replicates. Experimental groups were based on mouse phenotypes, and treatments were applied to primary cells isolated from individual mice per genotype per experiment. No statistical methods were used to predetermine sample sizes; they were based on experience with the methodology. For AFM measurements, outlier identification was carried out to remove rare individual measurements that represented apparent artefacts.

## Acknowledgements

We thank Karolina Punovuori and Satu-Marja Myllymäki for support with experiments, Hanne Ahola and Claudia Ortmeier for expert technical assistance, and the Max Planck Institute BioOptics and Biomedicum Imaging Unit, HiLIFE for support with imaging. This work was supported Doctoral Programme in Biomedicine (DPBM), Biomedicum Young Investigator grant (both to AJ), Instrumentarium Science Foundation and Academy of Finland postdoctoral fellowships (to AS), European Union’s Horizon 2020 research and innovation programme under Marie Skłodowska-Curie grant agreement no. 101032331 (to CV), Academy of Finland Research Fellowship 332821 (to LCB), the Sigrid Juselius Foundation, Helsinki Institute of Life Science, and Academy of Finland Center of Excellence BarrierForce and R’Life Programme consortium NucleoMech and the Max Planck Society (all to SAW).

## Author contributions

AJ designed and performed most of the experiments and analyzed data. AS and CV performed experiments and analyzed data. FP supported TEWL analyses. MR and CMN provided conceptual advice and supported establishment of assays. LCB supervised *in vivo* experiments and provided conceptual advice. SAW conceived and supervised the study, designed experiments and analyzed data. AJ and SAW wrote the paper. All authors read and commented on the manuscript.

## Supplementary Figure Legends

**Supplementary Figure 1.**
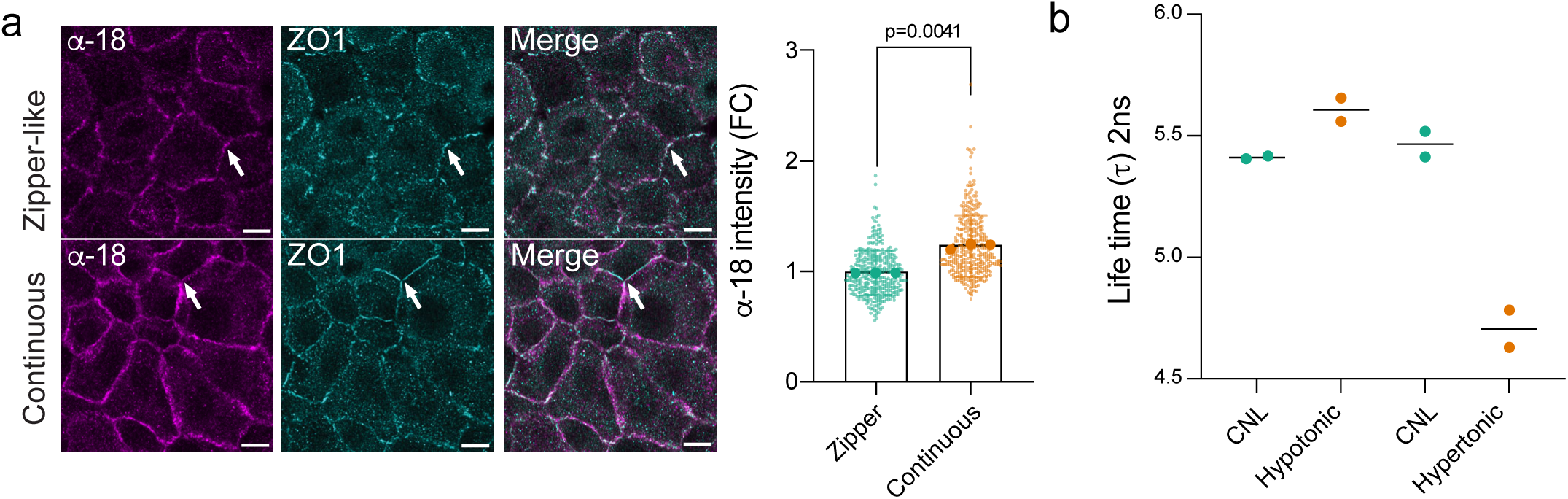
Analyses of junction and membrane tension after calcium switch or osmotic shocks. **a)** Representative images and quantification of tension-sensitive a-catenin (a-18) antibody and ZO1-stained primary keratinocytes fixed at 4 h or 8 h post Ca^2+^ addition. Arrows indicate zipper-like and continuous junctions. Note the increase of a-18 intensity at continuous junctions (n=3 independent experiments; mean±SD; Wilcoxon test; scale bar 10 µm). **b)** Quantification of fluorescence lifetime imaging of FLIPPER-TR from hypotonic and hypertonic controls from primary keratinocytes (n=2 independent experiments with >10 membrane measurements per condition/experiment).

**Supplementary Figure 2.**
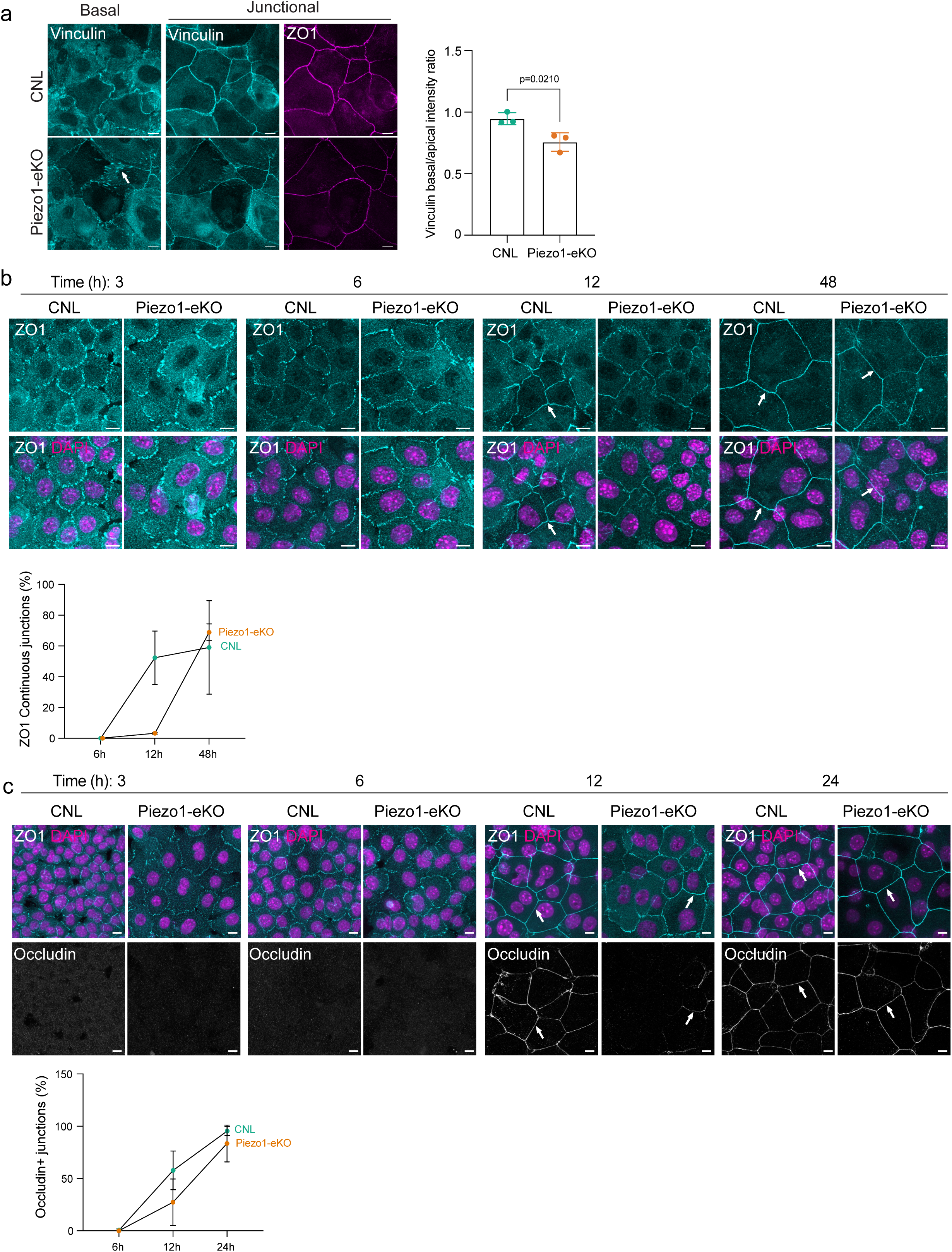
Piezo1-eKO cells show adhesion maturation defects. **a)** Representative images of vinculin- and ZO1-stained CNL and Piezo1-eKO primary keratinocytes fixed 24 h post Ca^2+^ switch. Quantification shows increased intensity of vinculin (arrow) in Piezo1-eKO basal focal adhesion plane compared to apical junctional plane (n =3 independent experiments; unpaired t-test; scale bars 10µm). **b)** ZO1- and DAPI-stained primary keratinocytes fixed at indicated time points post Ca^2+^ switch. Arrows show ZO1 in mature junctions of CNL and Piezo1-eKO cells at 12 h and at 48 h Quantification shows delayed junction maturation of ZO1 junctions in Piezo1-eKO cells (n=3 independent experiments; mean±SD; scale bar 10 µm). **c)** ZO1-, DAPI- and occludin-stained primary keratinocytes fixed at indicated time points post Ca^2+^ switch. Arrows indicate occludin-positive junctions. Quantification shows delayed emergence of occludin-positive tight junctions in Piezo1-eKO cells (n=3 independent experiments; mean±SD; scale bar 10 µm).

**Supplementary Figure 3.**
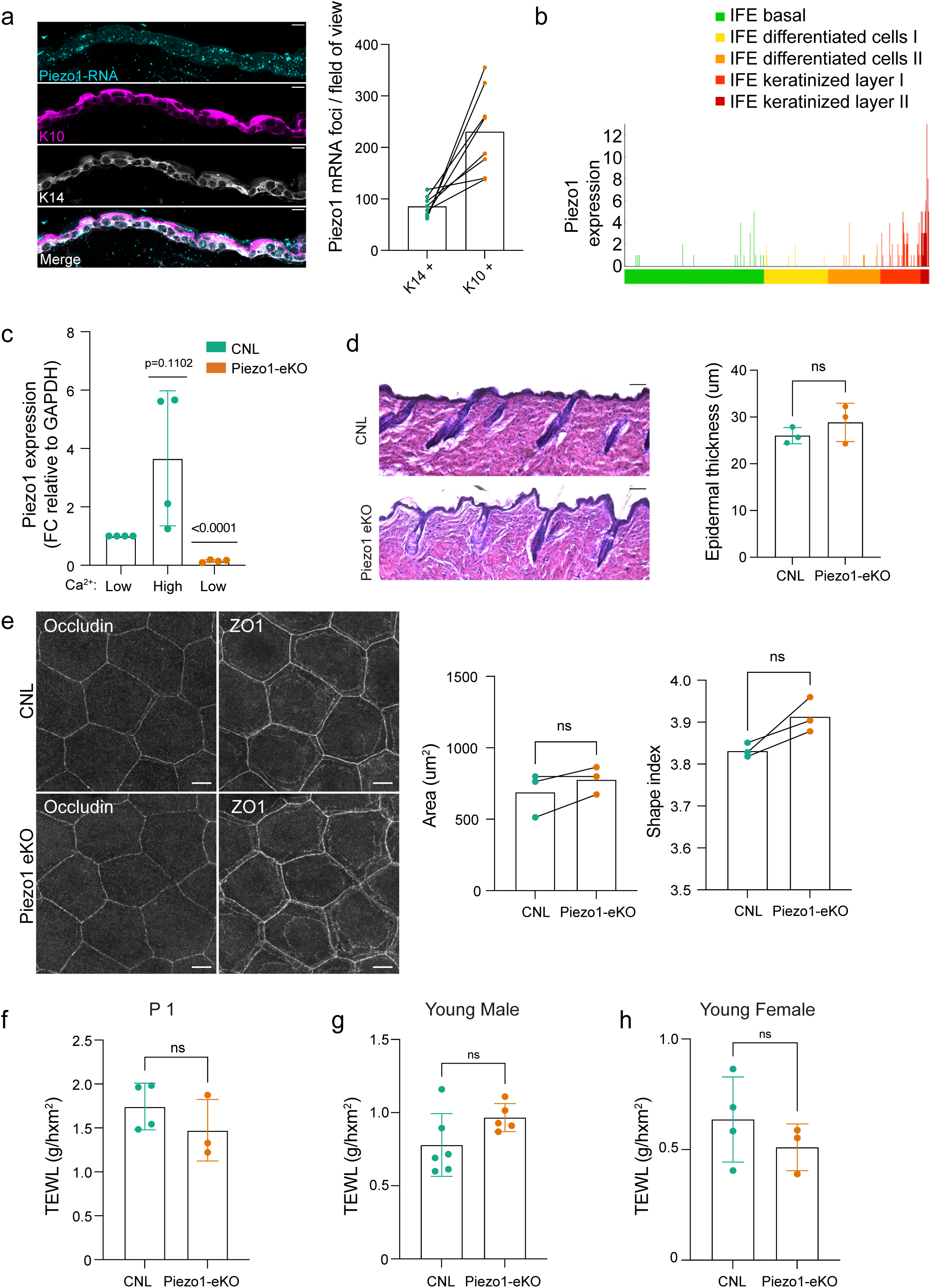
Analyses of Piezo1 expression and epidermal phenotype in young Piezo1-eKO mice. **a)** Representative images of 1-y old mouse back skin stained for Piezo1-RNA, Keratin-10 (K10) and Keratin-14 (K14) (n=8 fields of view). **b)** Analysis of Piezo1 mRNA expression from single cell RNAseq data from adult mouse back skin shows most abundant mRNA levels in suprabasal layers ^47^. **c)** RT-qPCR shows increased expression of Piezo1 in cells cultured under high Ca^2+^ calcium (n = 4 independent experiment, Kruskal-Wallis/Dunn’s). **d)** Representative H/E-stained back skin sections of CNL and Piezo1-eKO back skin mice (3-6 months old). Quantification comparable epidermal thickness between genotypes (n= 3 mice/genotype; unpaired t-test; scale bar 100 µm). **e)** Representative images and quantification of cell area and shape ZO1-stained ear whole mounts from 3-month-old CNL and Piezo1-eKO mice. (n =3 mice / genotype, paired t-test, scale bar 10 µm). **f)** Quantifications of transepidermal water loss measurements from back skin of CNL and Piezo1-eKO at postnatal day 1 (P1) (n=4 (CNL) and 3 (Piezo1-eKO) mice; unpaired t-test). **g)** Quantifications of transepidermal water loss measurements from back skin of 3 months old male CNL and Piezo1-eKO (n=6 (CNL) and 5(Piezo1-eKO) mice; unpaired t-test. **e)** Quantifications of transepidermal water loss measurements from back skin of 3 months old female CNL and Piezo1-eKO (n=4 (CNL) and 3 (Piezo1-eKO) mice; unpaired t-test). ns=not significant.

**Supplementary Figure 4.**
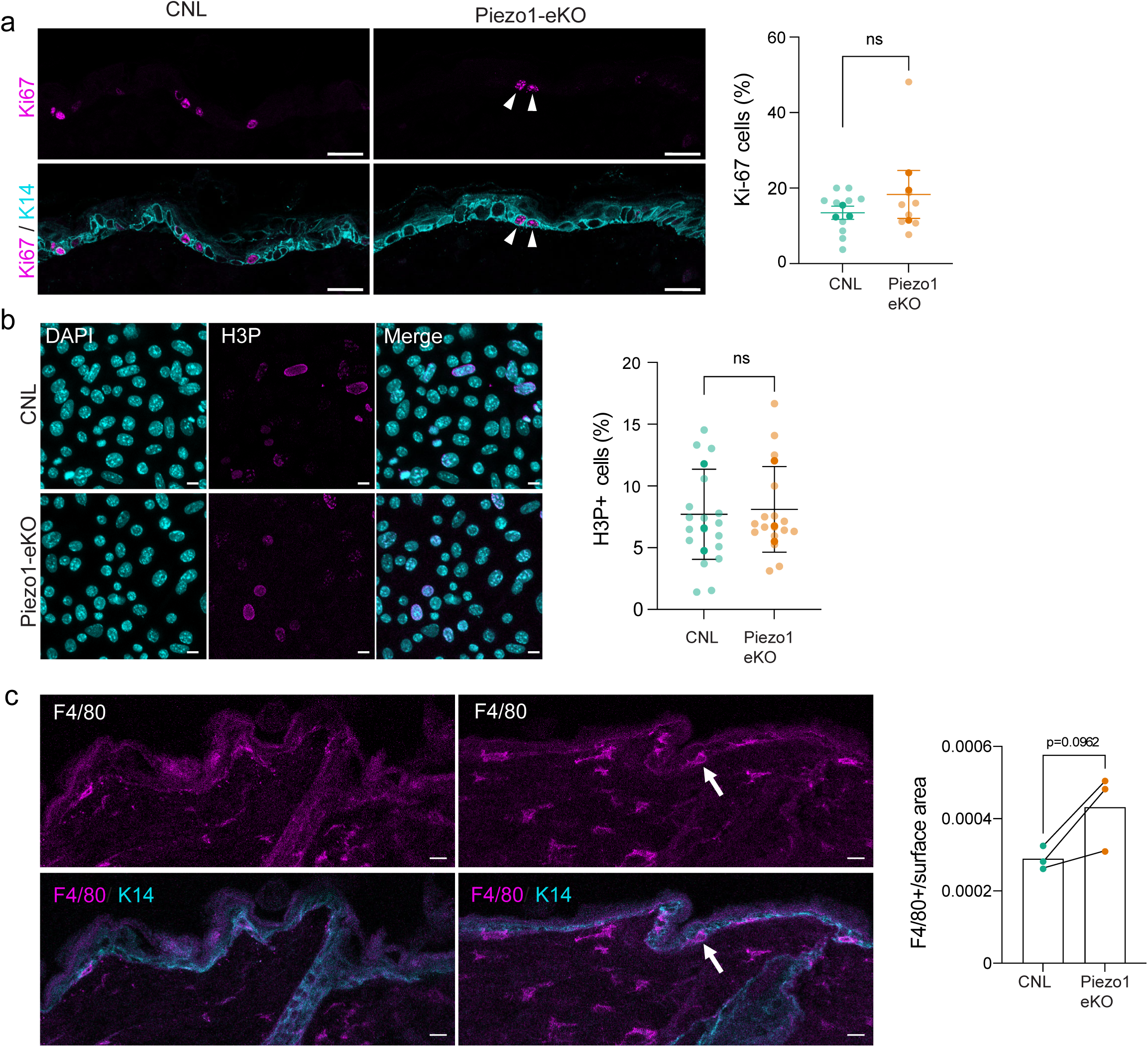
Analyses of epidermal phenotype of aged Piezo1-eKO mice. **a)** Representative images and quantification of CNL and Piezo1-eKO mice (young 3-6 months old) back skin stained for Ki-67 (magenta) and Keratin-14 (K14; cyan). Quantifications show no significant difference in proliferation (arrowheads) between CNL and Piezo1-eKO mice (n=3 mice / genotype, Mann-Whitney, scale bar 20 µm). **b)** Representative images and quantification of phosphorylated Histone H3 (H3P)-stained CNL and Piezo1-eKO cultured primary keratinocytes. No differences in proliferation were observed *in vitro* (n=3 independent experiments; paired t-test; scale bar 10 µm). **c)** Representative images and quantification of CNL and Piezo1-eKO <1-year old mice back skin stained for F4/80 (magenta) and Keratin-14 (K14; cyan). Note increased presence of F4/80 positive cells (arrows) in the skin of Piezo1-eKO mice (n=3 mice/genotype; paired t-test; scale bar 10 µm). ns= not significant.

**Supplementary Video 1**

**Analysis of adhesion maturation upon calcium addition in cultured primary keratinocytes**

Live imaging of keratinocytes transfected with ZO1-mEmerald. Note the transition of zipper-like adherens junctions into continuous “belt-like”, junctions starting around 5 h post Ca^2+^ switch. Arrow indicates the mature belt-like junction, asterisks mark untransfected neighboring cells. Acquisitions were performed at a rate of 20 min frame^−1^ for 16 h. Scale bars, 10 µm.

**Supplementary Video 2**

**Comparison of adhesion maturation between CNL and Piezo1-eKO keratinocytes**

Live imaging of *In vitro* cultured CNL and Piezo1-eKO keratinocytes transfected with ZO1-mEmerald. Note the faster transition of zipper-like junctions into continuous junctions in CNL cell (left) post Ca^2+^ switch. Arrow indicates the maturing mature belt-like junction, asterisks mark untransfected neighboring cells. Acquisitions were performed at a rate of 20 min frame^−1^ for 16 h. Scale bars, 10 µm.

